# The Impact on Estimations of Genetic Correlations of the Use, in Genome Wide Case-Control Studies, of Super-Normal, Unscreened and Family-History Screened Controls

**DOI:** 10.1101/693614

**Authors:** Kenneth S. Kendler, Chris Chatzinakos, Silviu-Alin Bacanu

## Abstract

Traditionally, in *normal* case-control studies of disorder A, individuals with disorder A are screened-out of controls. However, in genome wide association (GWA) studies, controls are sometimes *unscreened* or screened for disorder A and disorder B, producing *super-normal* controls. Using simulations, we examine how the observed genetic correlations between two disorders (A and B) are influenced by the use of unscreened, normal, and super-normal controls. Normal controls produce unbiased estimates of the genetic correlation. However, unscreened and super-normal controls both bias upward the genetic correlations. The strength of the bias increases with increasing population prevalences for the two disorders. With super-normal controls, the magnitude of bias is stronger when the true genetic correlation is low. The opposite is seen with the use of unscreened controls. Adding screening of first-degree relatives of controls substantially increases the bias in genetic correlations with super-normal controls but has minimal impact when normal controls are used.

---

Genetic correlations calculated from genome wide association (GWA) studies are widely used to quantify the genetic contribution to disorder comorbidity (Anttila et al., 2018; Bulik-Sullivan et al., 2015; Cross-Disorder Group of the Psychiatric Genomics Consortium(PGC-CDG), 2013). In this report, we examine one area of concern in such studies – that estimates of genetic correlations (r_g_) between disorders assessed using case-control GWA studies can be biased by the method of control selection.

In epidemiology, the proper selection of controls in case-control studies is a subject of considerable concern and debate (Hodge, Subaran, Weissman, & Fyer, 2012; Lubin & Gail, 1984; Wacholder, McLaughlin, Silverman, & Mandel, 1992; Wacholder, Silverman, McLaughlin, & Mandel, 1992a; Wacholder, Silverman, McLaughlin, & Mandel, 1992b). The principle defended by most methodologists is that controls should resemble cases in all characteristics *except* not having the disorder for which cases are selected – here disorder A. We term controls selected on this principle *normal* controls. Nonetheless, the collection of controls in many biomedical studies are neither properly done nor adequately reported (Lopez, Scheutz, Errboe, & Baelum, 2007; Malay & Chung, 2012).

In genetic epidemiology, some studies utilize *super-normal* controls screened not only for the disorder being studied but also for other often-related disorders (Chen et al., 2005; Schwartz & Susser, 2011). That is, if cases are selected for having disorder A, controls are selected for not having disorder A and for not having an additional screened-out disorder B (or multiple screened-out disorders B, C and D). In family studies, the use of super-normal controls produces spurious co-aggregation between disorders A and B the magnitude of which increases with increasing population prevalence of disorder B (Kendler, 1990). Case-control GWA studies have also utilized super-normal controls (Ikeda et al., 2011; Otowa et al., 2016; Schwartz & Susser, 2010; Sullivan et al., 2008).

Controls in case-control studies can be selected based not only on their own phenotype but also on the phenotype of close relatives (Wray et al., 2012). When normal controls or super-normal controls are screened in this fashion, we call these *normal-FH* and *super-normal-FH* controls, respectively, where FH stands for “family history.”

Finally, because screening of potential controls can be effortful and expensive, *unscreened* controls can be used (Kirov et al., 2009; O’Donovan et al., 2008). So, if cases are selected for having disorder A, investigators might take a population-based sample and not screen it so that it contains cases of A at approximately population prevalence.

Recently, van Rheenen et al. (van Rheenen, Peyrot, Schork, Lee, & Wray, 2019) provided a thorough review of the properties of GWA derived genetic correlations, finding the measure to be robust to many misspecifications of the causal model, case control proportions, and stratification. However, they provide two examples of what they call “double-screening control cohorts,” equivalent to what we have termed super normal controls. Consistent with results found in family studies (Kendler, 1990), they report minimal bias when the two disorders (schizophrenia and bipolar illness in their example) are relatively rare. However, when the two disorders are common (asthma and hay fever in their example), r_g_ between the two disorders is substantially inflated.

In this report, using simulation, we systematically explore the impact of different approaches to control selection in GWA studies on the estimation of r_g_, addressing four specific questions. First, as predicted by theory, does the use of normal controls in GWA studies produce unbiased estimates of r_g_? Second, attempting to replicate and expand upon the results presented by van Rheenen et al. (van Rheenen et al., 2019), do we find that the use of super-normal controls in GWA studies upwardly bias estimates of r_g_ between disorders A and B (r_gAB_)? If so, what is the effect on this bias of the true value of r_g_ and the population prevalences of A and B? Third, if we use normal-FH or super-normal-FH controls, does that contribute further to biases in r_gAB_? Finally, what bias occurs in r_gAB_ when the two disorders are studied using unscreened controls and how is that bias affected by the population prevalence of disorders A and B and the true r_g_ between them?

## METHODS

Due to the difficulty of performing theoretical calculations, we resorted to computationally intensive simulations using the Julia language version 0.6.4 (https://julialang.org/). For simplicity and computational tractability, we assumed that i) the studies of the two disorders of interest (A and B) are non-overlapping with total sample sizes ***N***_1_ = ***N***_2_ = **5,000**, for ii) GWAS consisted of ***M*** = **5,000** independent Single Nucleotide Polymorphisms (SNPs) and iii) super-normals are defined as screened out if they have either disorder A or B. The experiment assumes that the genetic correlation is computed, as it is commonly performed, from the GWAS summary statistics using equation (1) from Bulik-Sullivan et al (Bulik-Sullivan et al., 2015). Under the above settings for sample sizes and number of markers, the genetic covariance is simply ***E***(*z*_1*j*_*z*_2*j*_), where *z*_1*j*_ and *z*_2*j*_ are the ***j***-th SNP’s Z-score for the first and second disorder, respectively. Similarly, the heritabilities of the traits are 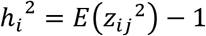, for ***i*** = **1,2**.

Let the two disorders under investigation have prevalences ***K*_1_** / ***K*_2_**, (equal) heritabilities 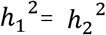 and genetic correlation ***ρ***. The genetic correlation matrix of the two disorders can be written as 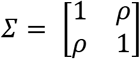. The varying parameters in the simulation design consisted of true genetic correlation, heritability and prevalence of the two disorders A and B (Table 1). **10** simulations were performed at each setting, such that the average standard error of the estimates was < **0.02**.

The simulation procedure is as follows:

1. Simulate (liability) random effects for the ***i***-th SNP (***i*** = **1,…,*M***) ***c_i_*~Bivariate Normal(0,*∑*)**.
2. Draw at random the minor allele frequency (MAF) of the ***i***-th SNP: ***p_i_*~Uniform(0.05,0.5)**.
3. Independently simulate the genotype for the father and the mother of the proband at each SNP. E.g., if the genotype at the ***i***-th SNP is ***G_i_*** = the number of minor alleles, then ***G_i_*~Bmomial(2, *p_i_*)**.
4. Simulate proband and sib by randomly sampling one allele from each parent
5. For each individual, assign the standard normal (Gaussian) liability for the ***j***-th disorder as 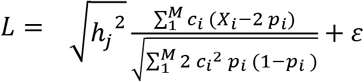 where 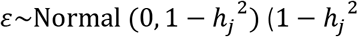 being the variance of the normal deviate).
6. For the ***j***-th trait assign case/control status for each member of the family when the upper percentile of the liability under a Gaussian distribution is less/bigger than ***K_j_***. (Assign SN status when a subject is a control for both disorders.)
7. number of cases and controls/SN controls equals 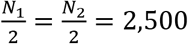.
8. Compute Z-score for the ***i***-th (***i*** = **1,2**) trait and ***j***-th SNP as 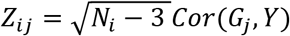, where ***G_j_*** is the genotype vector for cases and controls and ***Y*** is the phenotype vector (0 for controls and 1 for cases)

## RESULTS

We began by validating our simulation procedure by showing that for normal controls, the estimates of r_gAB_ are unbiased (Fig. 1, left panel). Prior to starting the simulation experiment, we also validated that the simulated liabilities for the two traits yield the desired genetic correlations (data not shown). We next examined the bias in r_gAB_ resulting from the use of super-normal controls with results depicted in figure 2, left panel. When both disorders A and B are rare (K1=K2=1.0%), the upward bias of r_gAB_ is very modest across the range of values of the true r_gAB_. When disorder K1 is rare (1.0%), and disorder K2 has a prevalence of 7.5% or 15%, the upward bias becomes more pronounced with greater bias when the true value of r_gAB_ is low. For example, when K2=7.5 and 15% and the true r_gAB_ equals 0.10, the estimates values (SE) of r_gAB_ are +0.18 (0.01) and +0.20 (0.01), respectively. The upward bias on r_gAB_ is slightly greater when both K1=K2=7.5% and becomes pronounced when the prevalences of K1 and K2 are 7.5 and 15%, and both are 15%. For example, if K1=K2=15%, when the true value of r_gAB_ equals 0, 0.10, 0.30 and 0.50, the estimated values of r_gAB_ equal, respectively, +0.31 (0.02), +0.40 (0.01), +0.54 (0.01), and +0.68 (0.01).

**Figure 1.**
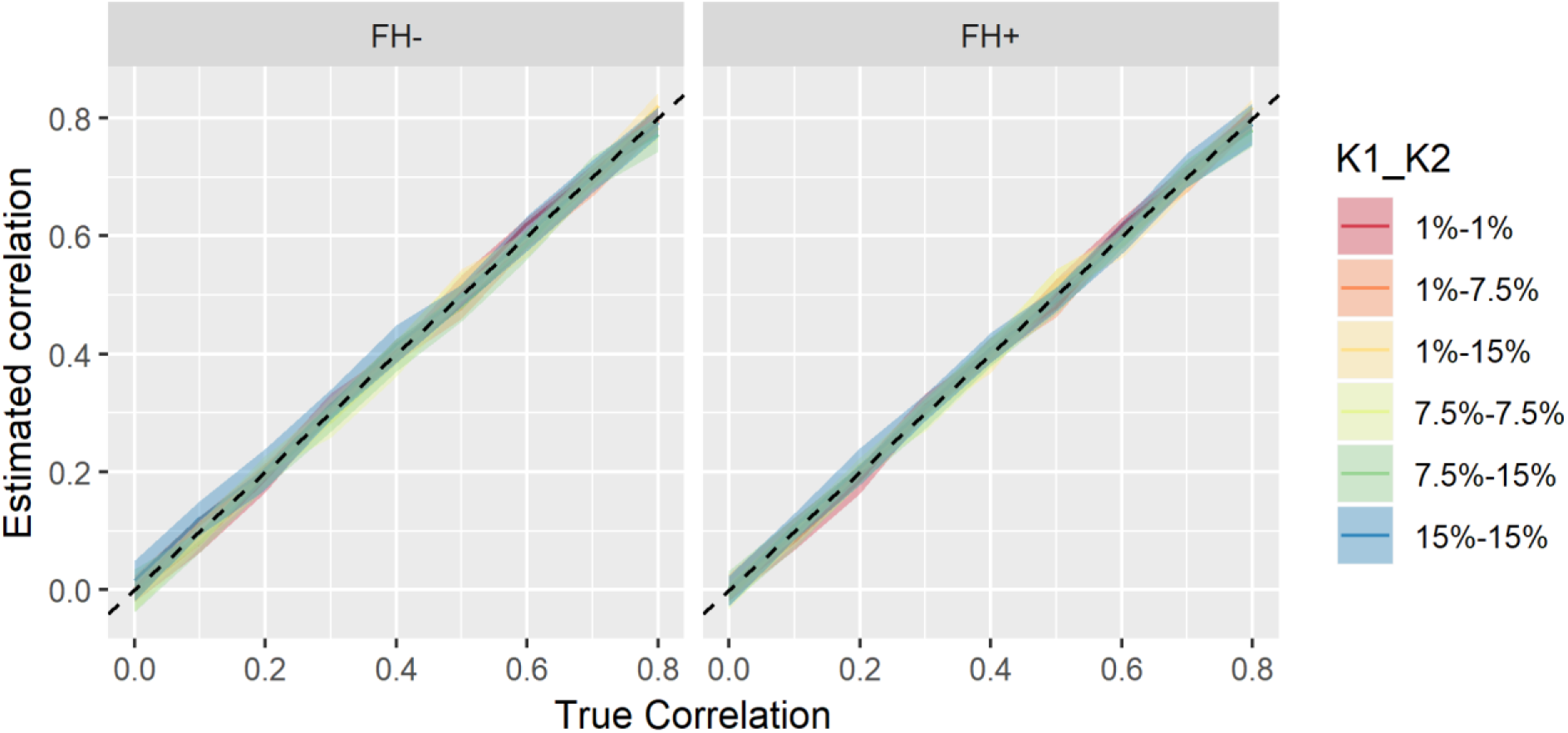
The relationship between the estimated correlation (as-per-Bulik-Sullivan et al. (Bulik-Sullivan et al., 2015)) and true correlation and true correlation when both disorders use normal controls. The banner settings relate to family history (FH), with (FH+) or without (FH-) family history screening for the condition(s). Each disorder is assumed to have a population prevalence of 1%, 7.5% and 15%, the color of different lines defines the assumed prevalence pair of the disorders traits (K1_K2-K1 for disorder A and K2 for disorder B). The heritability of each trait was assumed to be 0.5. The black dotted line denotes an unbiased estimator, i.e. estimated and true correlations are equal.

We then examined the effects of using normal-FH and super-normal-FH controls. As seen in figure 1 right panel, using normal-FH controls produced no appreciable bias in the estimated r_gAB_. However, compared to standard super-normal control, super-normal-FH controls produced a greater upward bias on r_gAB_ (figure 2, right panel) which was particularly strong when the true value of r_g_ between the two disorders equaled zero. For example, when K1=K2=1.0%, K1=K2=7.5% and K1=K2=15.0%, the estimated values of r_gAB_ equaled, respectively, +0.04 (0.01), +0.27 (0.01), and +0.50 (0.01).

**Figure 2.**
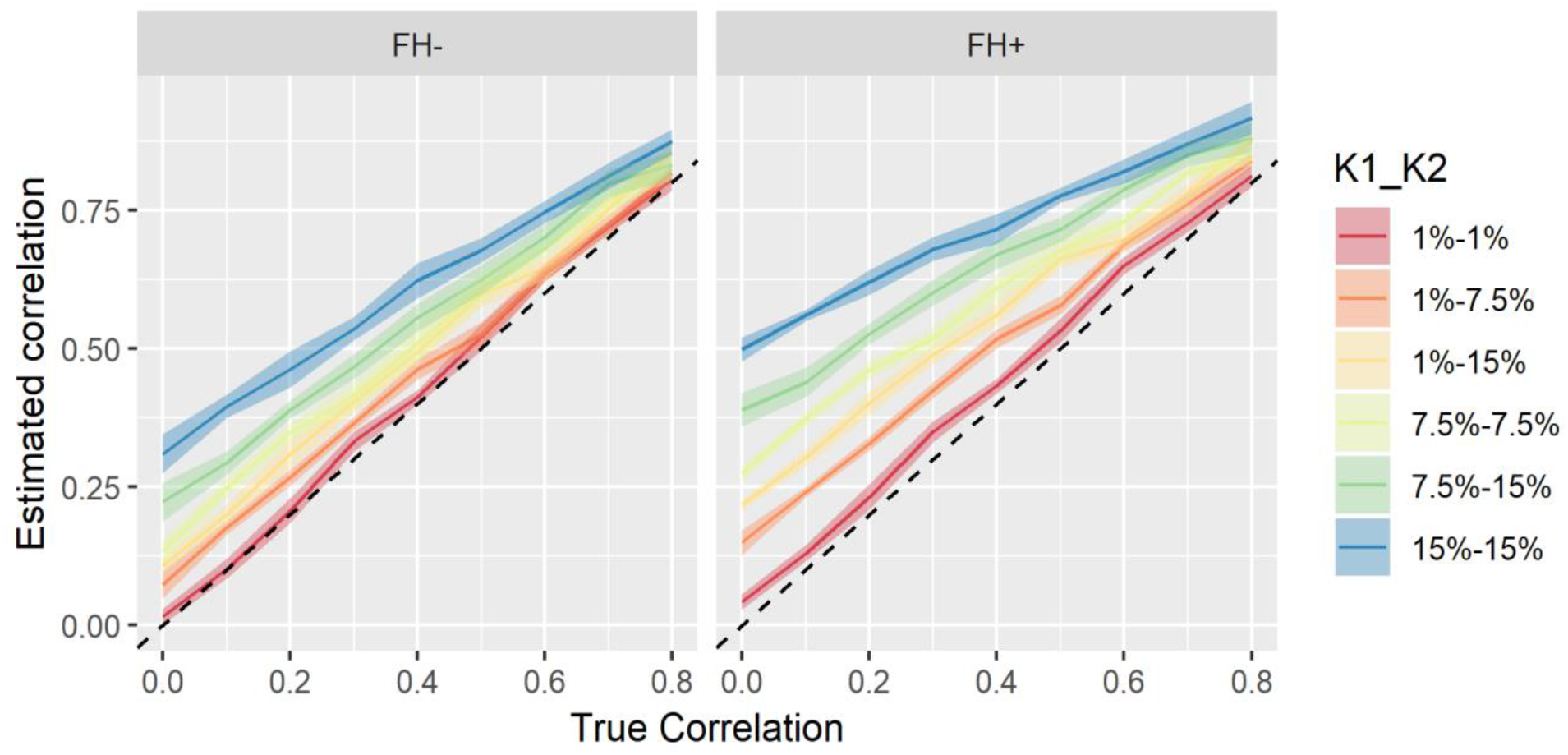
The relationship between the estimated correlation and true correlation when both disorders are studied using-super-normal controls as a function of the true correlation and the prevalence rates of the two disorders (K1 for disorder A and K2 for disorder B). For notation and background see Fig.1.

Finally, we examined the impact of unscreened controls on estimates of r_gAB_ (figure 3). At low levels of true genetic correlation, the upward biases were modest. However, these biases became progressively stronger with higher true genetic correlations and higher prevalence disorders producing, in several situations, implausible values of genetic correlations exceeding unity.

**Figure 3.**
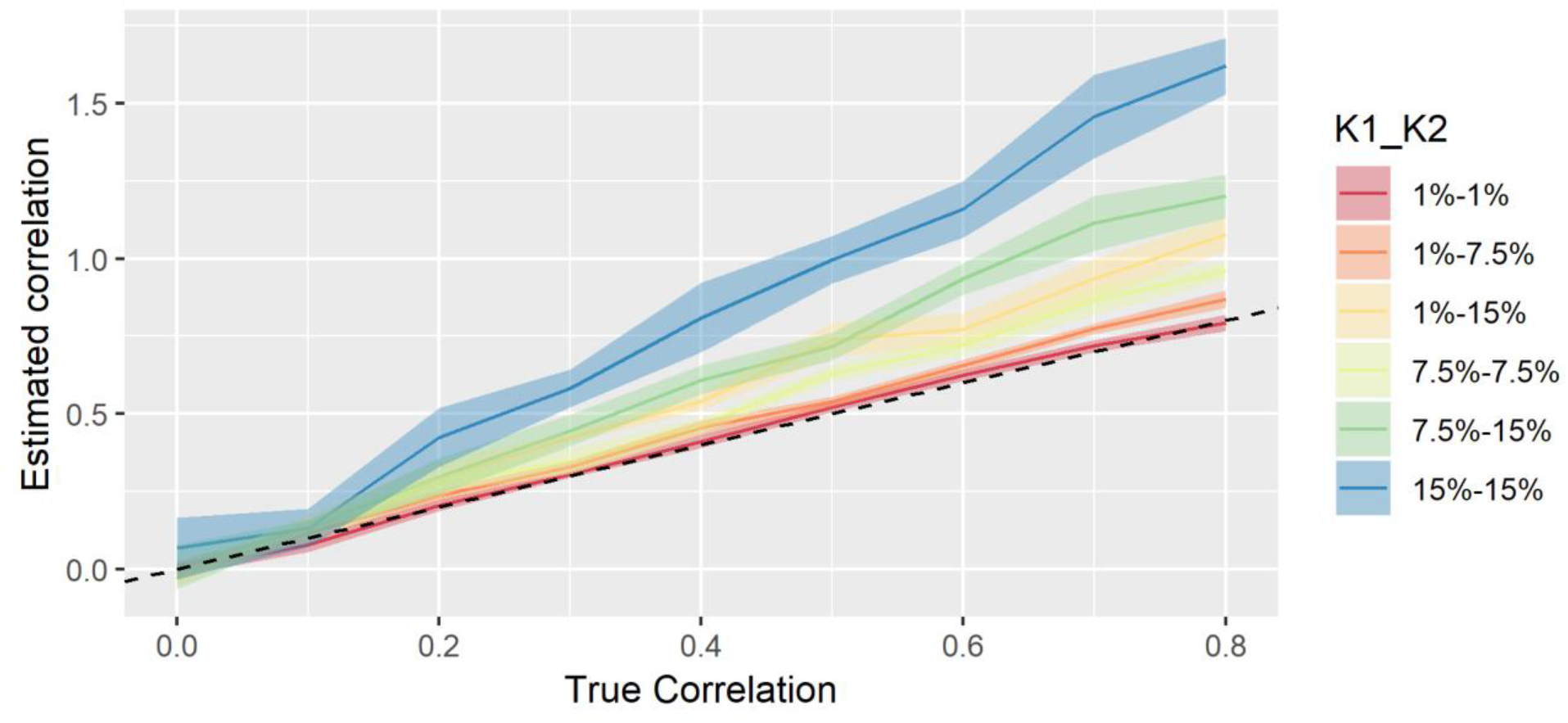
The relationship between the estimated correlation (as-per-Bulik-Sullivan et al (Bulik-Sullivan et al., 2015)) and true correlation when both traits use unscreened controls. For notation and background see Fig.1.

## DISCUSSION

We draw four major conclusions from these simulations. First, to obtain accurate estimates of r_g_ using GWA results, studies should use normal controls. Second, replicating and extending the results of van Rheenen et al. (van Rheenen et al., 2019), we find that using super-normal controls will result in upward biases the magnitude of which are strongly related to the prevalence of the pair of disorders and their true r_g_. At low prevalences (≤ 1%), the bias introduced is modest while it can become quite substantial at high prevalences (≥ 15%). The upward bias on estimate of r_g_ is also substantially stronger at low true correlations and, with high prevalence disorders, can produce evidence for substantial r_g_s where none exists. The replication of the results for super-normal controls is reassuring given that van Rheenen et al (van Rheenen et al., 2019) derived their results theoretically, while we used a simulation design coupled with empirical estimation of correlation (Bulik-Sullivan et al., 2015) in order to explore other types of controls (e.g. unscreened or screened together with close relatives).

Third, while screening the close relatives of controls produced no appreciable bias in r_g_ when used with normal controls, screening relatives using a super-normal control paradigm substantially exacerbated the upward bias on r_g_s. Finally, while the use of unscreened controls in GWA studies can result in savings of research resources, they pose a risk of substantially upwardly biasing evidence for genetic correlations especially when used with relatively common disorders.

These results are intuitively plausible. Using normal controls, the signal in a case-control GWA study arises entirely from the fact that all the cases and none of the controls have disorder A. When disorder B is eliminated from only controls, the case-control difference becomes “contaminated” and reflects not only case-control differences in the prevalence of A but also of B. The same will occur when we study disorder B but screen our controls for disorder A. So, if we took the results of such a GWA study and calculated r_gAB_, our estimates would be biased upward because our genetic risk for disorder A is contaminated with risk variants for disorder B and vice-versa. Within this thought experiment, it is obvious that the strength of the “contamination” of our GWA study will relate directly to the frequency of the screened-out disorder. If rare, it will impact only modestly on the resulting “signal.” If we screen relatives of controls for disorder B when studying A, this will further reduce the genetic risk in controls for disorder B, increasing the difference with cases and contaminating the signal and hence the resulting r_gAB_. On the other hand, unscreened controls induce an increase in genetic correlation due to mainly underestimating the heritability of the trait(s). That is, the heritability is underestimated due to the average noncentrality of statistics being lower when compared to their screened control counterparts (see genetic covariance and heritability estimates in Appendix).

These results should be interpreted in the context of three methodological limitations. First, we only presented results at a single heritability for our two disorders: 0.50. We repeated results for a higher heritability (0.80 – see appendix). The pattern of results was similar. Second, we only presented results where the same control selection method had been used with both disorders. In real life, r_g_ can be calculated between one disorder assessed using normal controls and another using super-normal controls. Simulations suggested that, in this case, the results were generally in between those found when normal and super-normal controls were assessed for both diseases. Third, we have not considered an expanded super-normal control method that screens for multiple disorders. Here results would depend on the prevalence and true genetic correlations between the range of disorders excluded from controls and would likely produce stronger and more wide-spread biases than presented here when we examined only two disorders at a time.

## Supplementary Material

### Estimates for genetic correlation results for trait heritability of 0.8

When compared to a heritability of 0.5 for both traits, the results are rather similar to their homologues (Fig. S1–3). The one slight exception is that the bias is now diminished when studies use population (Fig. S1). These biases are still sizeable, especially when i) not using normal controls and ii) one of the traits is very common.

**Figure S1.**
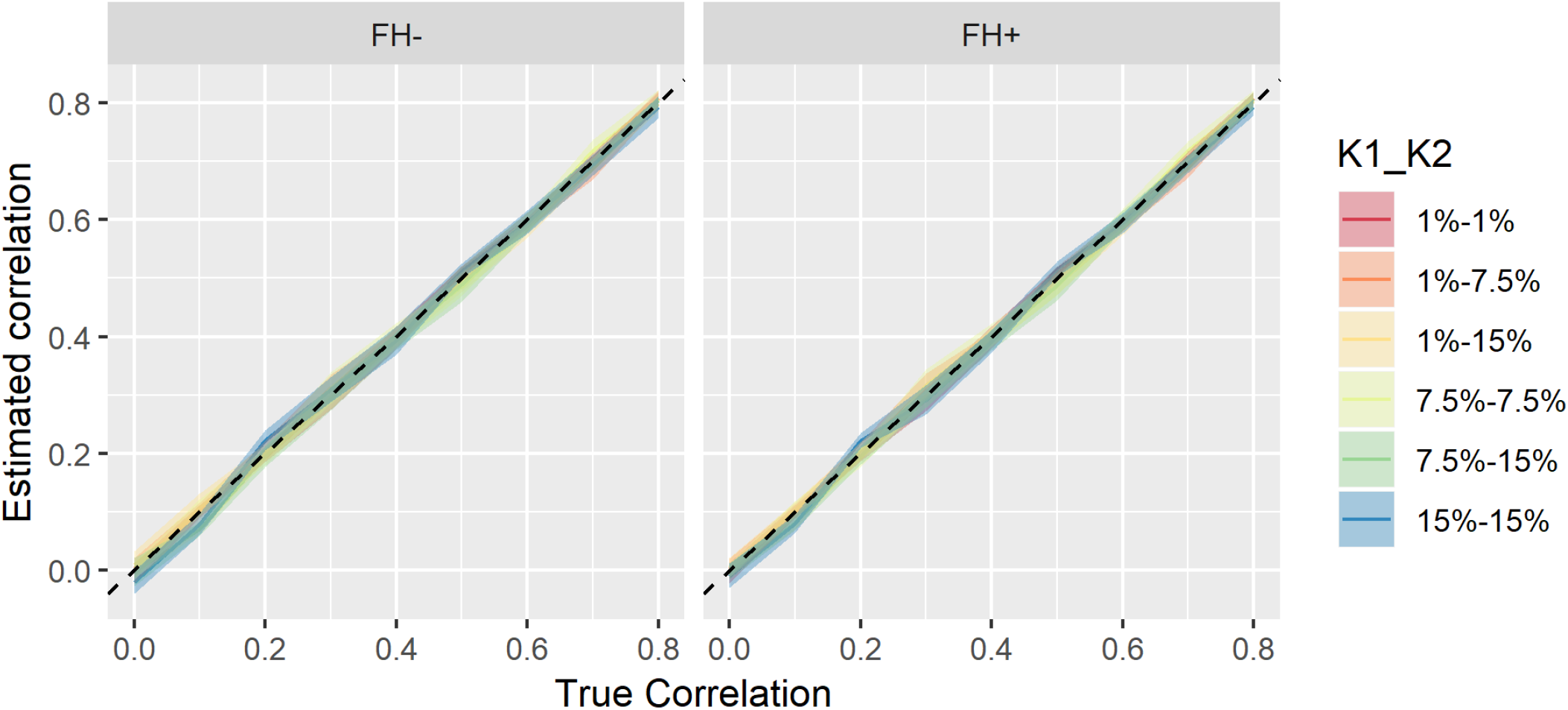
The relationship between the estimated correlation and true correlation and true correlation when both traits use normal controls. The banner settings relate to family history (FH), with (FH+) or without (FH-) family history screening for the condition(s). Each trait is assumed to have prevalence of 1%, 7.5% and 15%, the color of different lines defines the assumed prevalence pair of the traits (K1_K2-where K1 for disorder A and K2 for disorder B). The heritability of each trait was assumed to be 0.8. The black dotted line denotes an unbiased estimator, i.e. estimated and true correlations are equal.

**Figure S2.**
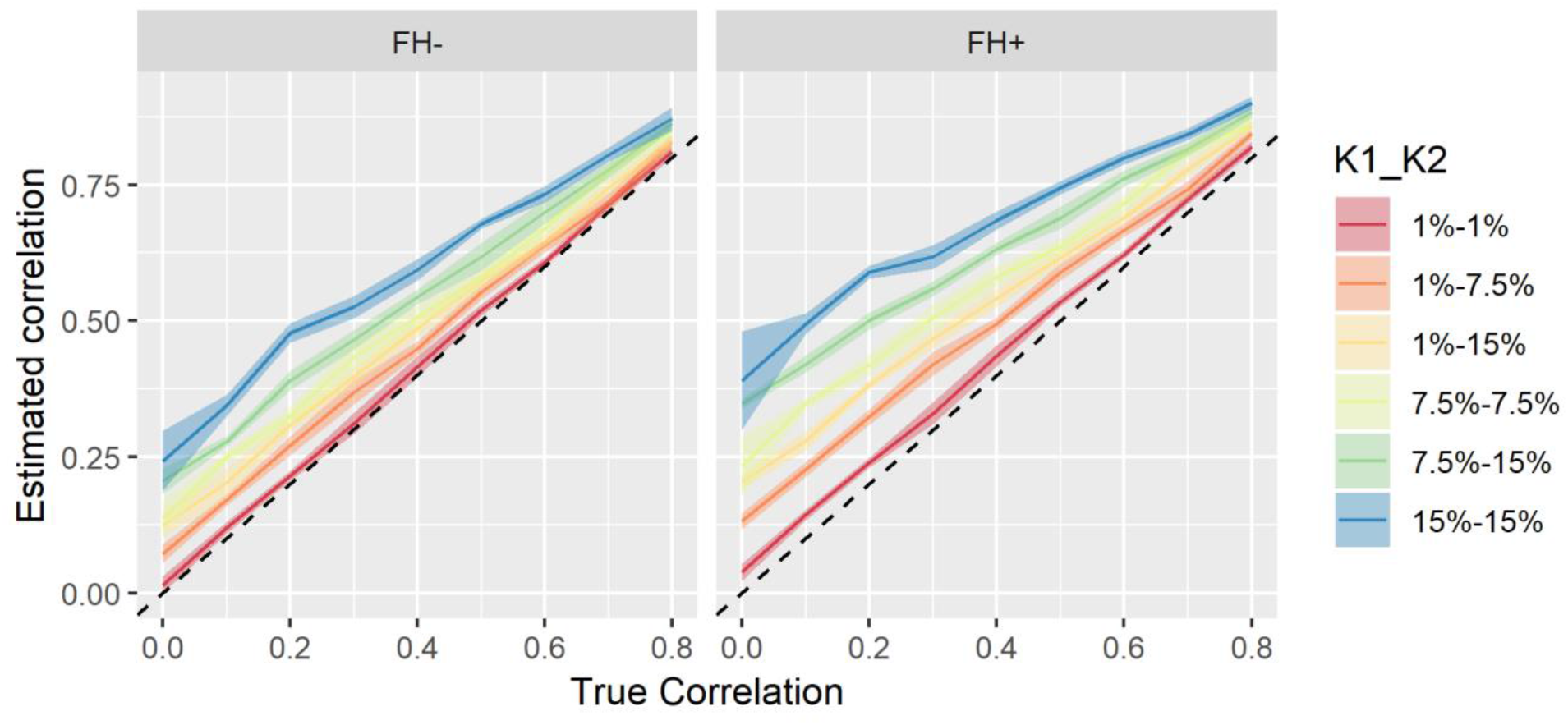
The relationship between the estimated correlation and true correlation and true correlation when both traits use super-normal controls low prevalence traits. For notation and background see Fig.S1.

**Figure S3.**
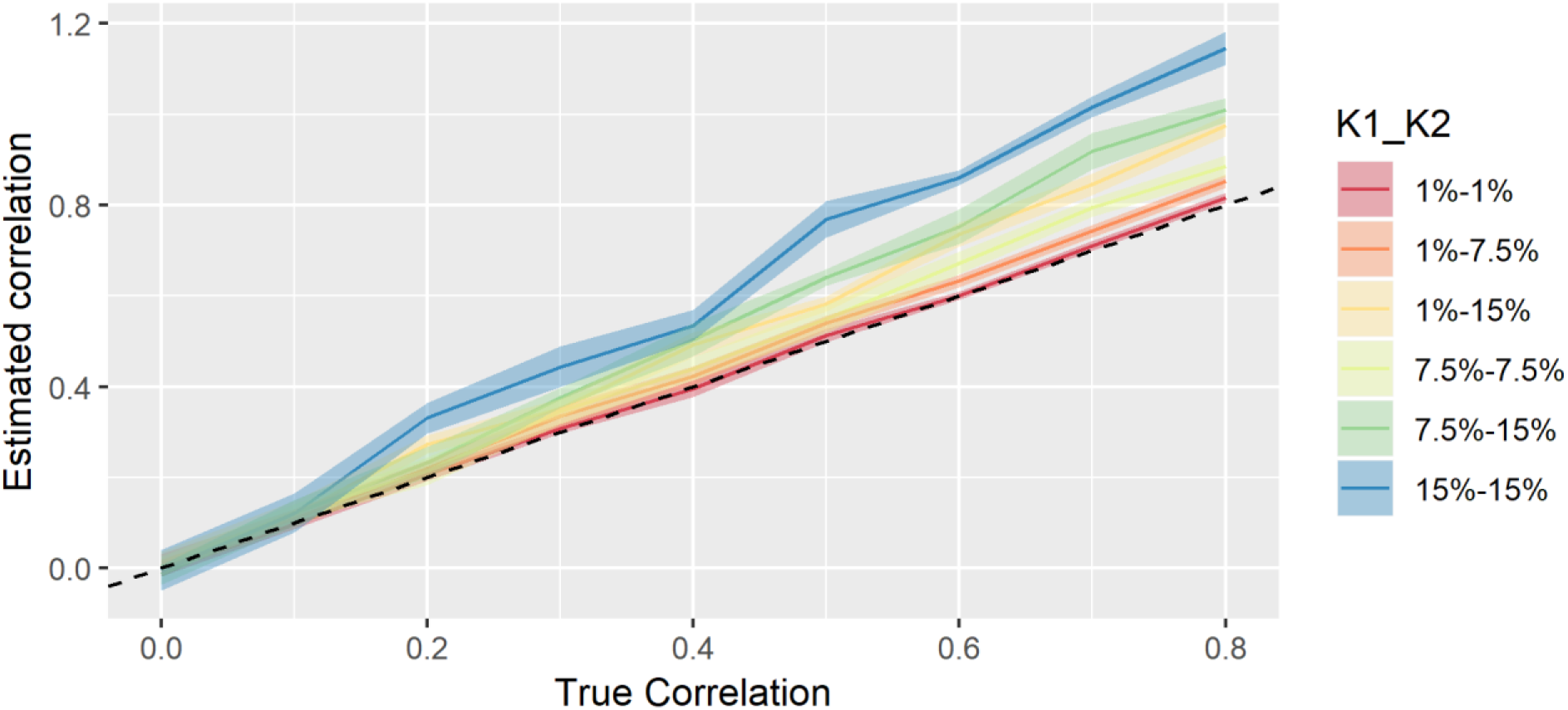
The relationship between the estimated correlation (as-per-Bulik-Sullivan et al.^1^) and true correlation when both traits use unscreened controls.. For notation and background see Fig.S1.

### Components of genetic correlation estimation

The genetic correlation is the quotient of genetic covariance and the square root of heritabilities. It is, thus, of interest to assess how the estimates of these two measures vary for unscreened/super-normal controls relative to the normal controls. Consequently, we also estimated heritabilities and genetic covariances under the settings presented in the main manuscript.

The main points relating to genetic covariance estimates are: i) at high prevalences, the use of normal controls induce a slight overestimation only under the FH scenario and population controls a modest underestimation and ii) super-normal controls induce large overestimations, especially for lower true genetic correlations and/or super-normal-FH (Fig. S3–S6). The main feature for estimated heritabilities are: i) at high prevalences, the use of normal controls induces a modest overestimation (only under FH scenario) and population controls a sizeable underestimation ii) super-normal controls induce reasonably large overestimations, especially for lower true genetic correlations and/or super-normal-FH controls (Fig. S7–S9).

**Figure S4.**
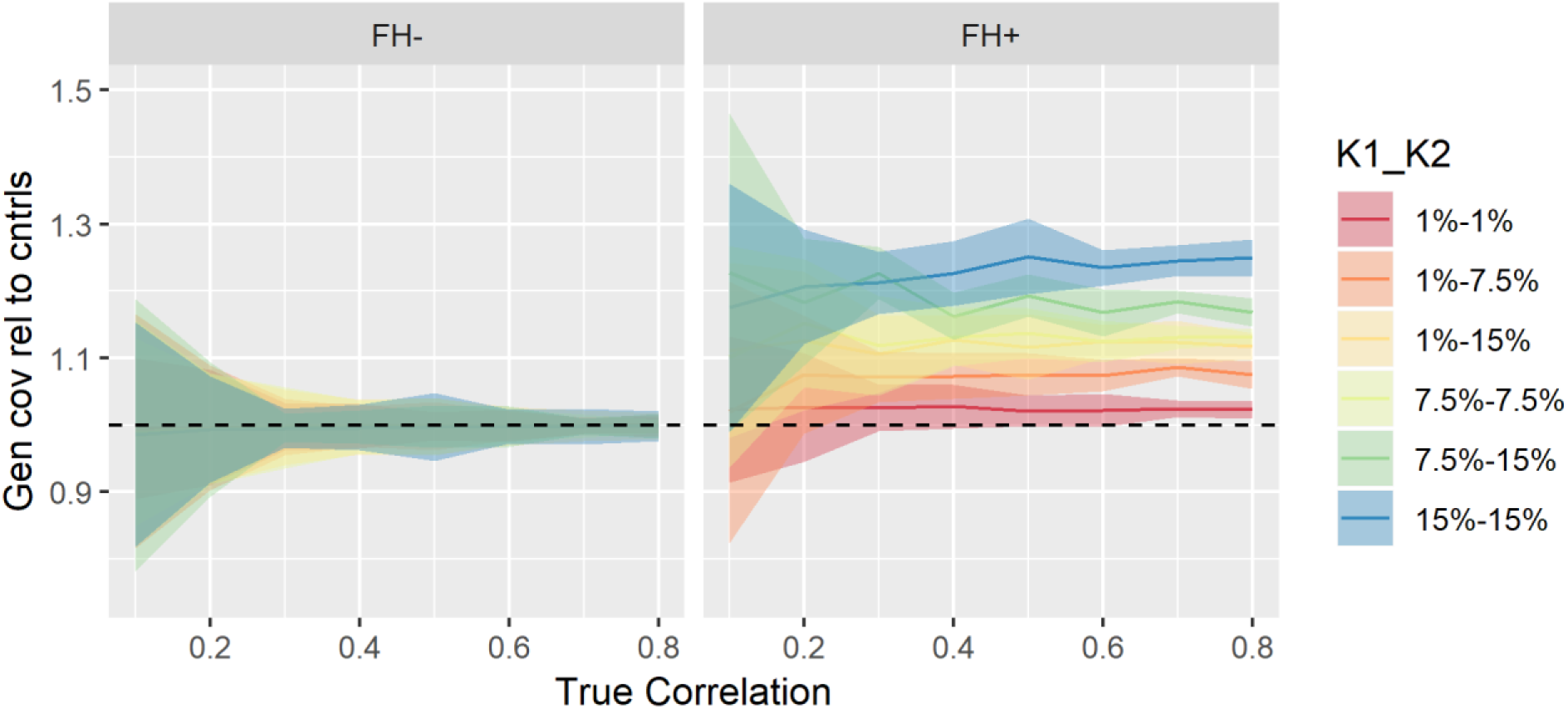
The relationship between the estimated genetic covariance (relative to normal controls) and true correlation when both traits use normal controls. The heritability of each trait was assumed to be 0.8. The true correlation value of 0 was eliminated from the plot due to the division with an estimate of zero (estimated genetic covariance of studies with normal controls), which resulted in extremely large confidence intervals. The black dotted line denotes an unbiased estimation in normal controls (i.e. the horizontal line at y=1). For notation and background see Fig.S1.

**Figure S5.**
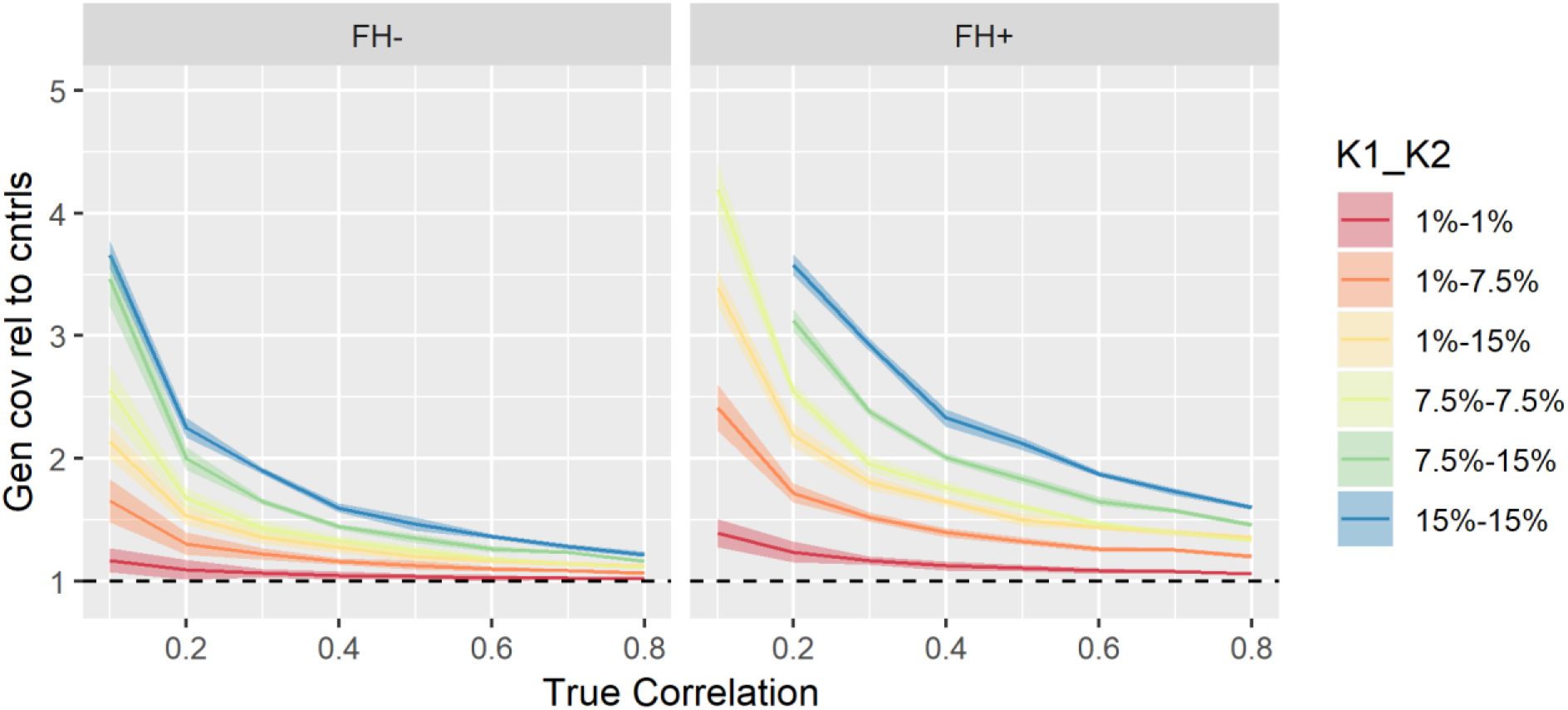
The relationship between the estimated genetic covariance (relative to normal controls) and true genetic correlation when both traits use super-normal controls. The heritability of each trait was assumed to be 0.8. For notation and background see Fig.S1 and S4.

**Figure S6.**
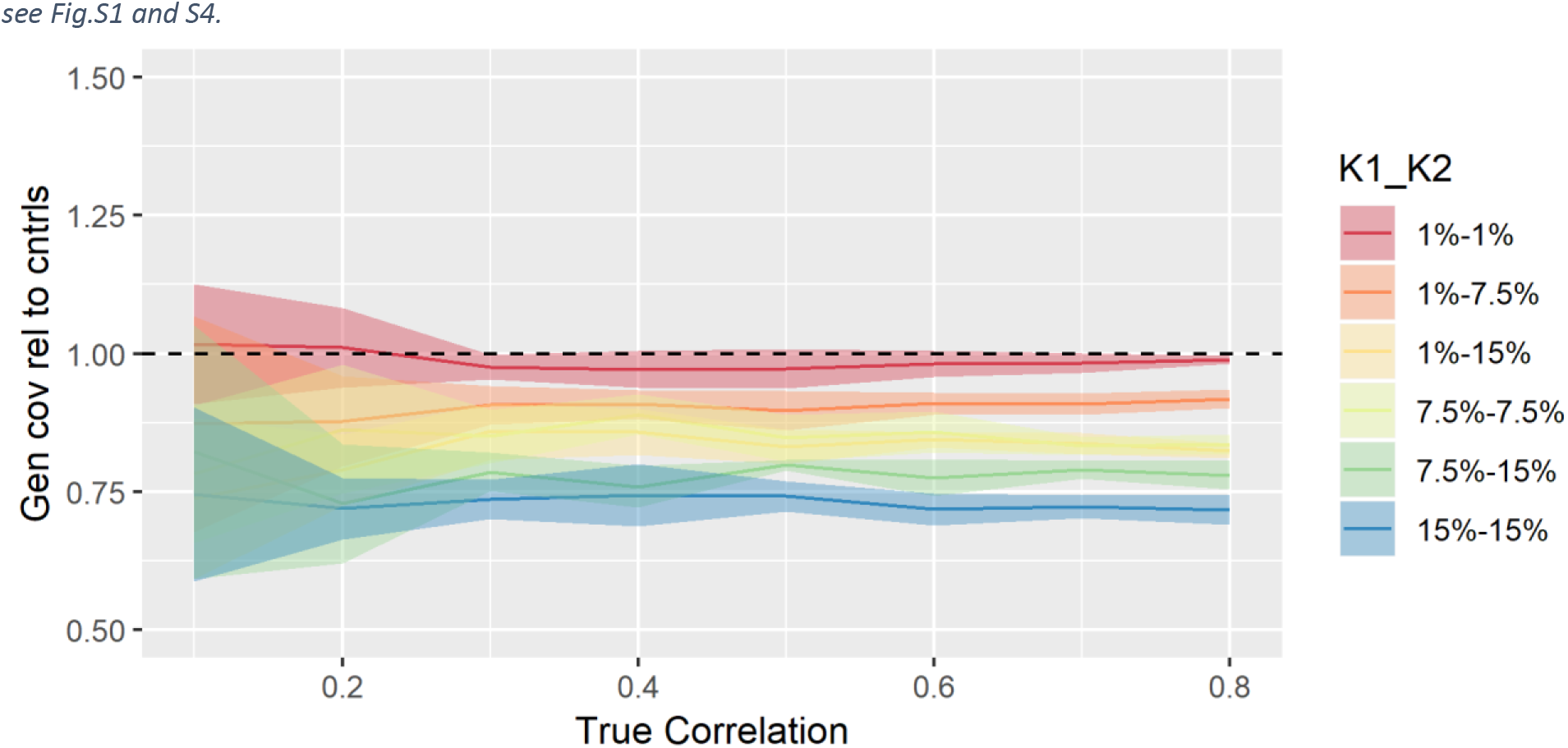
The relationship between the estimated genetic covariance (relative to normal controls) and true genetic correlation when both traits use unscreened controls. The heritability of each trait was assumed to be 0.8. For notation and background see Fig.S1 and S4.

**Figure S7.**
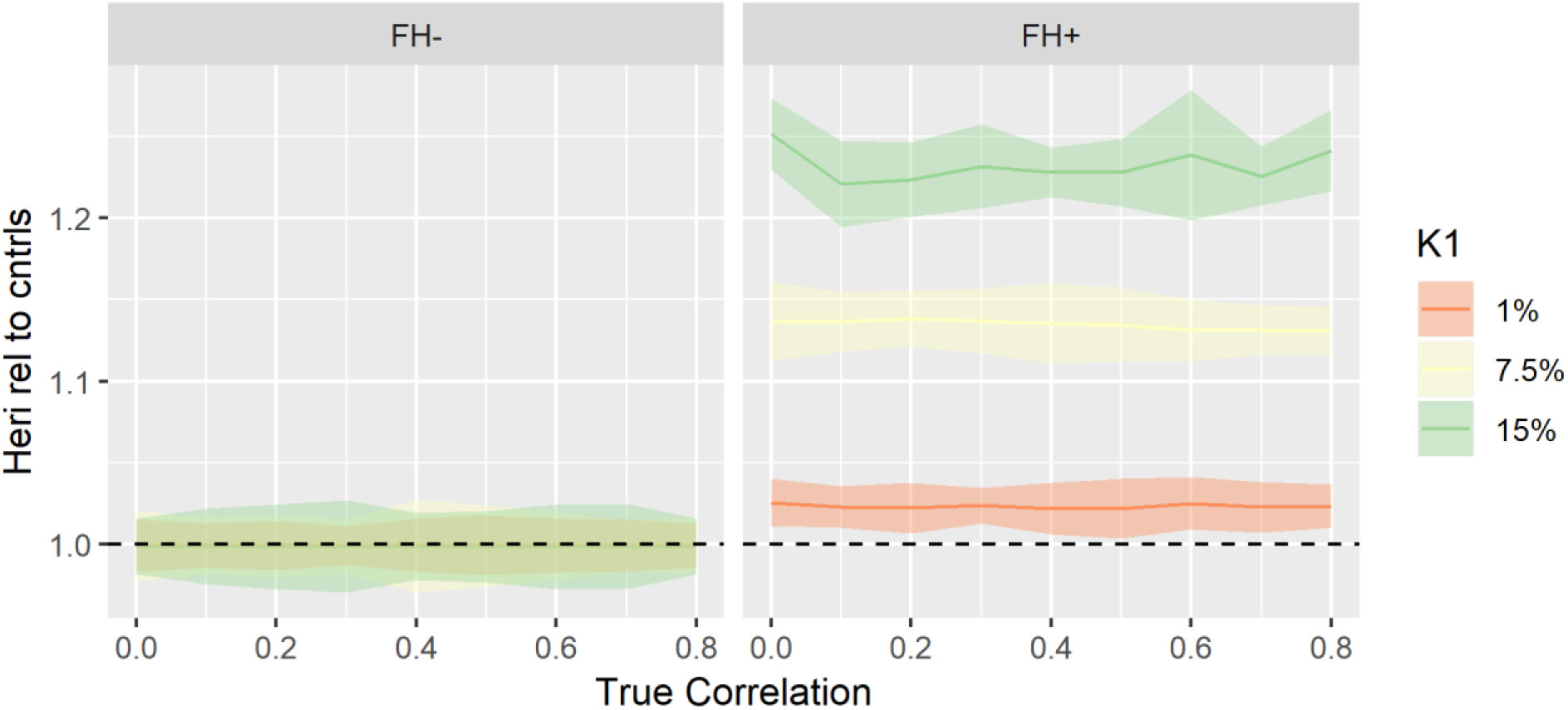
The relationship between the estimate in heritability (relative to normal controls) and true correlation when both traits use normal controls. The black dotted line denotes an unbiased estimation in normal controls (i.e. the horizontal line at y=1). For notation and background see Fig.S1 and S4.

**Figure S8.**
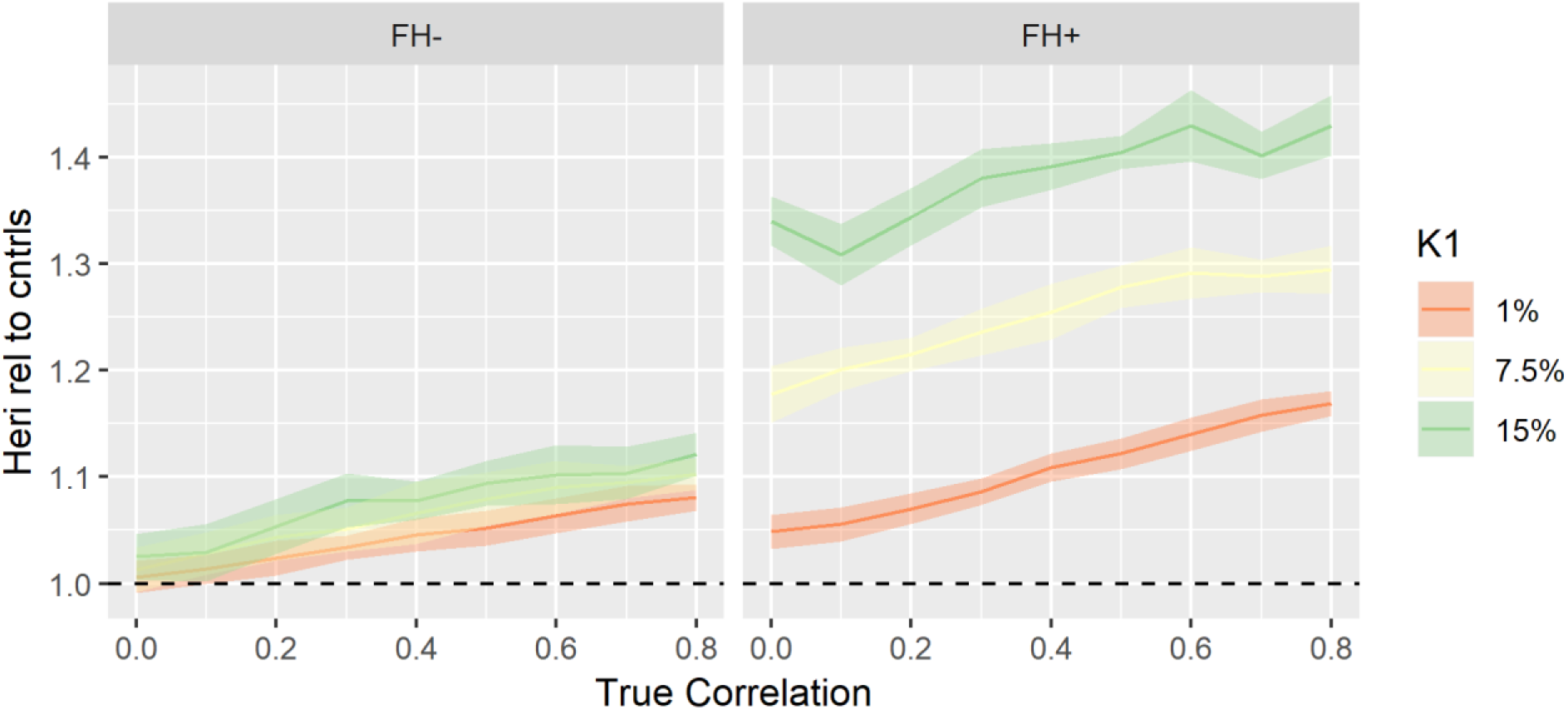
The relationship between the estimated heritability (relative to normal controls) and true genetic correlation when both traits use super-normal controls. The heritability of each trait was assumed to be 0.8. For notation and background see Fig.S1 and S7.

**Figure S9.**
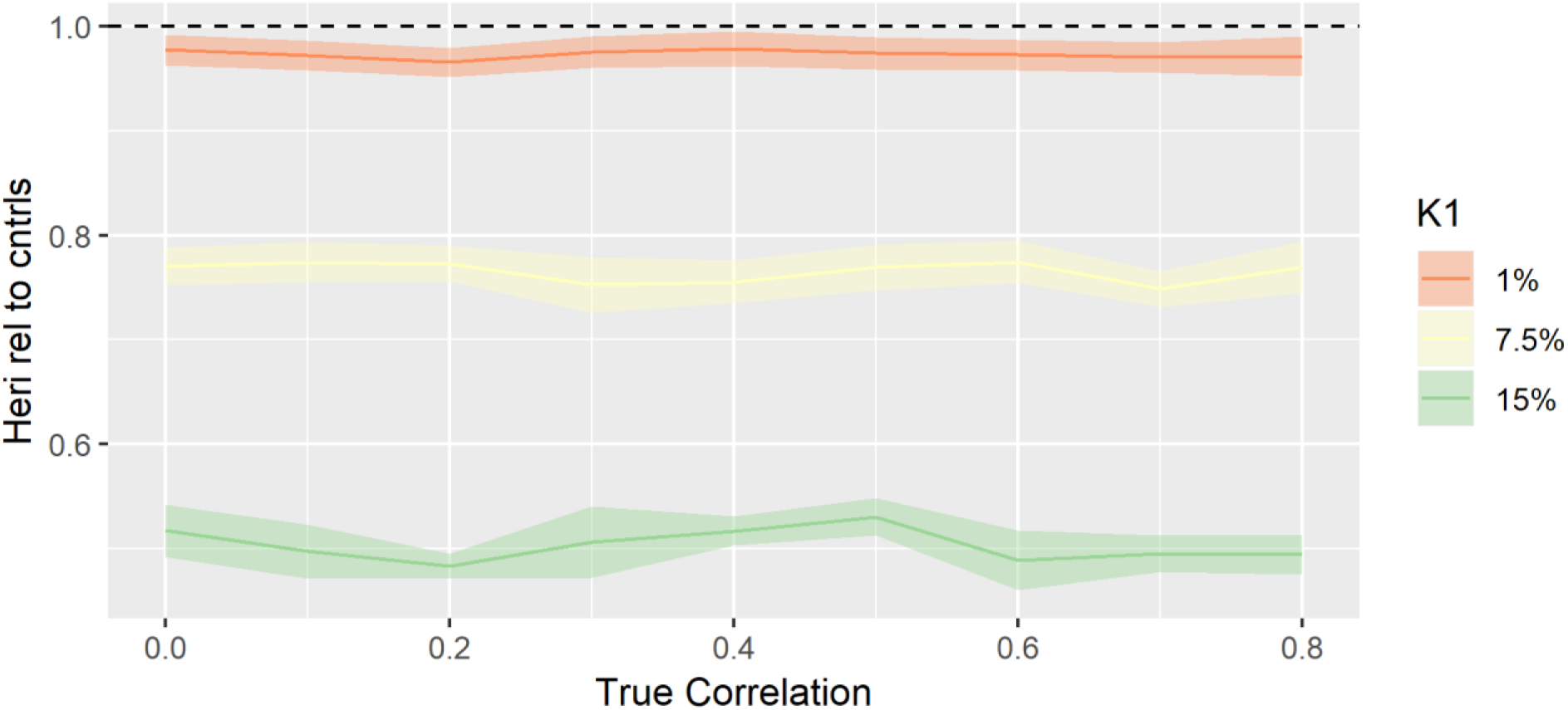
The relationship between the estimated heritability (relative to normal controls) and true genetic correlation when both traits use unscreened controls. The heritability of each trait was assumed to be 0.8. For notation and background see Fig.S1 and S7.

